# TREM2 Orchestrates Myeloid Cell Programming and Immune Dysregulation in Pulmonary Hypertension

**DOI:** 10.64898/2026.09.18.752808

**Authors:** Aline C. Oliveira, Filipe da Silva, Yutao Zhang, Matthew D. Alves, Ann T. Pham, Charlotte Harris, Sultan Khanfar, Rafael Abdelnor, Jimena Alvarez-Castanon, Caroline Phillips, Mia T. Hall, Chunhua Fu, Shiza T. Virk, Katherine e. Ray, Li Chen, Annette de Kloet, Eric Krause, Andrew J. Bryant

**Author notes:** **Corresponding Author**: Aline C. Oliveira, Department of Pharmacology and Therapeutics, 1200 Newell Drive| Gainesville, Fl 32610-0274, P.O. Box 100267|.

## Abstract

Background: Pulmonary hypertension (PH) is a progressive and fatal disease characterized by pulmonary vascular remodeling, inflammation, and immune dysregulation. Myeloid-derived suppressor cells (MDSCs) and macrophages contribute to PAH pathobiology; however, the molecular regulators of their pathological activation remain poorly defined. The triggering receptor expressed on myeloid cells 2 (TREM2) is an immunomodulatory receptor that shapes myeloid cell metabolism, survival, and immunosuppressive function, yet its role in PAH has not been investigated. Methods: Single-cell RNA sequencing (scRNA-seq) data from human pulmonary artery tissue (GSE210248; n=3 PAH, n=3 donors) were analyzed to characterize TREM2 expression across cell populations. Wild-type (WT) and global TREM2 knockout (TREM2 KO) mice were exposed to chronic hypoxia (10% FiO2, 28 days). Hemodynamic, histological, flow cytometric, and ex vivo functional assessments were performed. TREM2 protein expression was measured by flow cytometry in circulating MDSCs from PAH patients (n=22) and healthy controls (n=13). Results: TREM2 was markedly enriched in monocyte/macrophage (mono/macs) populations in PAH pulmonary arteries, with co-upregulation of APOE, GPNMB, and CSF1R. TREM2-high immune cells exhibited transcriptional downregulation of chemotaxis programs and upregulation of antigen processing and MHC II presentation pathways. TREM2 deficiency significantly attenuated hypoxia-induced RVSP increase, right ventricular dysfunction, pulmonary inflammation, and vascular remodeling. TREM2 KO mice showed blunted mTOR/p-S6 activation in bone marrow myeloid populations and MDSCs from TREM2 KO mice exerted significantly less suppression of CD4+ and CD8+ T cell proliferation. TREM2 KO mice tended to preserve bone marrow CD4+ T cell proportions and significant upregulation of CD62L on T cells under hypoxia. TREM2 was significantly elevated in circulating MDSCs from PAH patients and showed a directional association with hemodynamic severity. Conclusions: TREM2 is a novel regulator of myeloid-driven immunosuppression and vascular remodeling in PAH and warrants investigation as a therapeutic target and biomarker.

## INTRODUCTION

Pulmonary hypertension (PH) is a progressive and life-threatening condition characterized by elevated pulmonary arterial pressure (mPAP) that ultimately leads to right ventricular heart failure and significant morbidity and mortality.^1–4^ It is a rare but progressive disorder with an estimated prevalence of around 1% in the general population and up to 10% in those over 65 years of age.^3,5^ The natural history of patients with pulmonary arterial hypertension results in median survival times of 2.8 years following diagnosis.^6^ Despite advances in diagnosis and therapy, recent registry data suggest that, with modern treatments, median survival has improved, with 1-year, 3-year, and 5-year survival rates now estimated at approximately 85–90%, 70–75%, and 60–70% respectively.^3,6,7^ Nevertheless, PH continues to substantially shorten life expectancy, with marked detereioration in quality-of-life, particularly in patients with advanced disease or significant right ventricular dysfunction.

The underlying PH pathophysiology involves vascular remodeling, inflammation, and endothelial dysfunction,^1,8,9^ but the molecular mechanisms driving these processes remain incompletely understood. Recent transcriptomic and proteomic studies have highlighted the involvement of innate immune pathways in the vascular remodeling associated with PH,^10–13^ suggesting that immune modulators play a critical role in disease progression. In this context, PH is now recognized as not merely a vascular disease, but one with substantial immune and inflammatory involvement.^9,14^ Central to this immune response are myeloid cells, including macrophages and myeloid-derived suppressor cells (MDSCs), which serve as key mediators of tissue injury and repair.^15,16^ During pulmonary vascular remodeling, these cells can accumulate within the lung vasculature and undergo phenotypic shifts in response to local microenvironmental cues, thereby influencing inflammation, fibrosis, and vascular cell proliferation.^13,17^ Emerging evidence indicates that targeting regulatory pathways governing myeloid cell responses can modulate these processes and alter disease progression.^15–18^

Among the molecular regulators of myeloid cell function, the triggering receptor expressed on myeloid cells 2 (TREM2) has emerged as a key immunomodulatory receptor expressed predominantly on macrophages and other myeloid lineage cells.^19–21^ TREM2 functions as a lipid– and damage-associated sensor that shapes myeloid cell activation, metabolism, and survival in response to tissue stress.^19–21^ Through these mechanisms, TREM2 influences macrophage phenotypic adaptation in chronic inflammatory environments, regulating processes such as immune suppression, tissue remodeling, and fibrosis.^22–24^ While TREM2 signaling has been associated with protective roles in certain contexts, including maintenance of myeloid homeostasis, dysregulated or sustained TREM2 activation has been linked to pathological immunosuppression and impaired immune surveillance in chronic disease states.^22^ Notably, although TREM2 has been extensively studied in neurodegenerative and metabolic disorders, its role in cardiopulmonary disease, and particularly in pulmonary arterial hypertension, where chronic inflammation, hypoxia, and tissue remodeling are central features, remains poorly defined.

Given the essential contribution of the immune system, particularly myeloid populations such as macrophages and myeloid-derived suppressor cells (MDSCs), to pulmonary vascular remodeling, and the emerging recognition of TREM2 as a central regulator of myeloid responses in cancer and chronic inflammatory diseases, we hypothesized that TREM2 plays a critical role in the development of PH. To test this hypothesis, we combined analysis of human pulmonary vascular transcriptomic data with genetic loss-of-function approaches in a murine model of hypoxia-induced pulmonary hypertension to enable assessment of the contribution of TREM2 to cardiopulmonary remodeling under chronic hypoxic exposure.

## METHODS

### Single-Cell RNA Sequencing Data Analysis

To characterize TREM2 expression in the pulmonary vasculature of PAH patients, we re-analyzed a publicly available scRNA-seq dataset (GSE210248) derived from human pulmonary artery tissue of PAH patients and healthy donors (n=3 per group). Bioinformatics processing, cell-type annotation, GO enrichment analyses and more details of the materials and methods are provided in the Supplemental Material.

### Animals

Ten-to-twelve-week-old C57BL/6J WT and TREM2 KO (C57BL/6J-Trem2em2Adiuj/JStrain #:027197) male and female mice (n=6–10 per group) were exposed to normoxia (FiO₂ 21%) or chronic hypoxia (FiO₂ 10%) for 28 days in a normobaric ventilated chamber (Coy Laboratory Products). All procedures were approved by the University of Florida Institutional Animal Care and Use Committee (protocol number: IACUC202400000401).

### Hemodynamic Assessment

Under Avertin anesthesia (16 mg/kg intraperitoneally), RVSP and ±dP/dt were recorded using a 1.4-pressure-volume catheter (Millar Instruments, SPR-671) inserted via the right jugular vein as previously described.^25^ Right ventricular hypertrophy was quantified as the Fulton index [RV/(LV+S) weight ratio].

### Histological Analysis

Paraffin-embedded lung sections were stained with Masson’s trichrome and α-SMA. Pulmonary inflammation was graded on a validated 0–4 scale in a blinded manner across 10 random fields (10× magnification). Muscularized vessel counts and medial wall thickness were assessed by α-SMA staining.

### Flow Cytometry

Single-cell suspensions from bone marrow and lungs were stained with fluorochrome-conjugated antibodies and Fixable Viability Dye (eBioscience) for surface and intracellular targets and acquired on a BD FACSymphony A3 cytometer. The complete antibody panel and acquisition protocols are provided in Table S1 available in the Supplemental Methods.

### T Cell Suppression Assay

MDSCs were isolated from spleens of WT and TREM2 KO mice using the EasySep™ Mouse MDSC Isolation Kit (StemCell Technologies, 19867). T cells were isolated from spleens of naïve WT C57BL/6 mice using the EasySep™ Mouse T Cell Isolation Kit (StemCell Technologies, 19851), labeled with CellTrace™ Violet (Thermo Fisher), and co-cultured with MDSCs at MDSC:T cell ratios of 4:1, 2:1, and 1:1 for 5 days with anti-CD3/CD28 stimulation. Percent suppression was calculated as [1 − (% proliferation with MDSCs / % proliferation without MDSCs)] × 100.

### Human Samples

Peripheral Blood Mononuclear Cells (PBMCs) were isolated from pulmonary arterial hypertension (PAH) patients (n=22) and healthy controls (n=13) enrolled under IRB protocol number IRB201400744. PBMCs were differentiated in GM-CSF– supplemented medium for 7 days and assessed for TREM2 expression by flow cytometry. TREM2 levels were correlated with mPAP, PASP, and RVSP.

### Statistical Analysis

Data are presented as mean±SEM. Differences between groups were assessed by 2-way ANOVA with Tukey post-hoc correction. Where variances were substantially unequal, Welch unpaired 2-tailed t test was applied. All analyses were performed using GraphPad Prism 11. A P value <0.05 was considered statistically significant.

## RESULTS

### TREM2 expression is enriched in monocyte/macrophages populations in the pulmonary arteries of patients with PAH

TREM2 expression on peripheral macrophages modulates T-cell activation in inflammatory-mediated diseases such as cancer and atherosclerosis. Given that our previously published work on MDSC findings suggest similar immune-suppressive mechanisms in PH^25^, we investigated the role of TREM2 in its pathogenesis. To establish the clinical relevance of TREM2 expression in patients with PH, we analyzed a publicly available scRNA-seq dataset GSE210248 derived from pulmonary arterioles of donor controls and PAH patients (n=3 per group). Unsupervised clustering identified all major pulmonary cell populations, including fibroblasts, smooth muscle cells, endothelial cells, epithelial cells, granulocytes, mast cells, dendritic cells, NK cells, T/NK cells, B cells, and mono/macs clusters, across samples. Visualization of these populations, separated by condition, and analysis of cellular composition revealed shifts in the abundance of vascular and immune cell populations between donor and PAH samples (Figure 1A and B), consistent with an inflammatory and remodeling environment characteristic of PH. Cell type proportions show a fundamental compositional difference between PAH and donor pulmonary artery tissue (Figure S1). In PAH, T/NK cells constituted the single largest population, followed by NK cells, Mono/Macs, granulocytes, and endothelial cells. Feature plot visualization of TREM2 on UMAP embeddings for donor and PAH tissue separately confirmed these patterns further and demonstrated enhanced TREM2 expression within the PAH dataset (Figure 1B; Figure S1).

**Figure 1.**
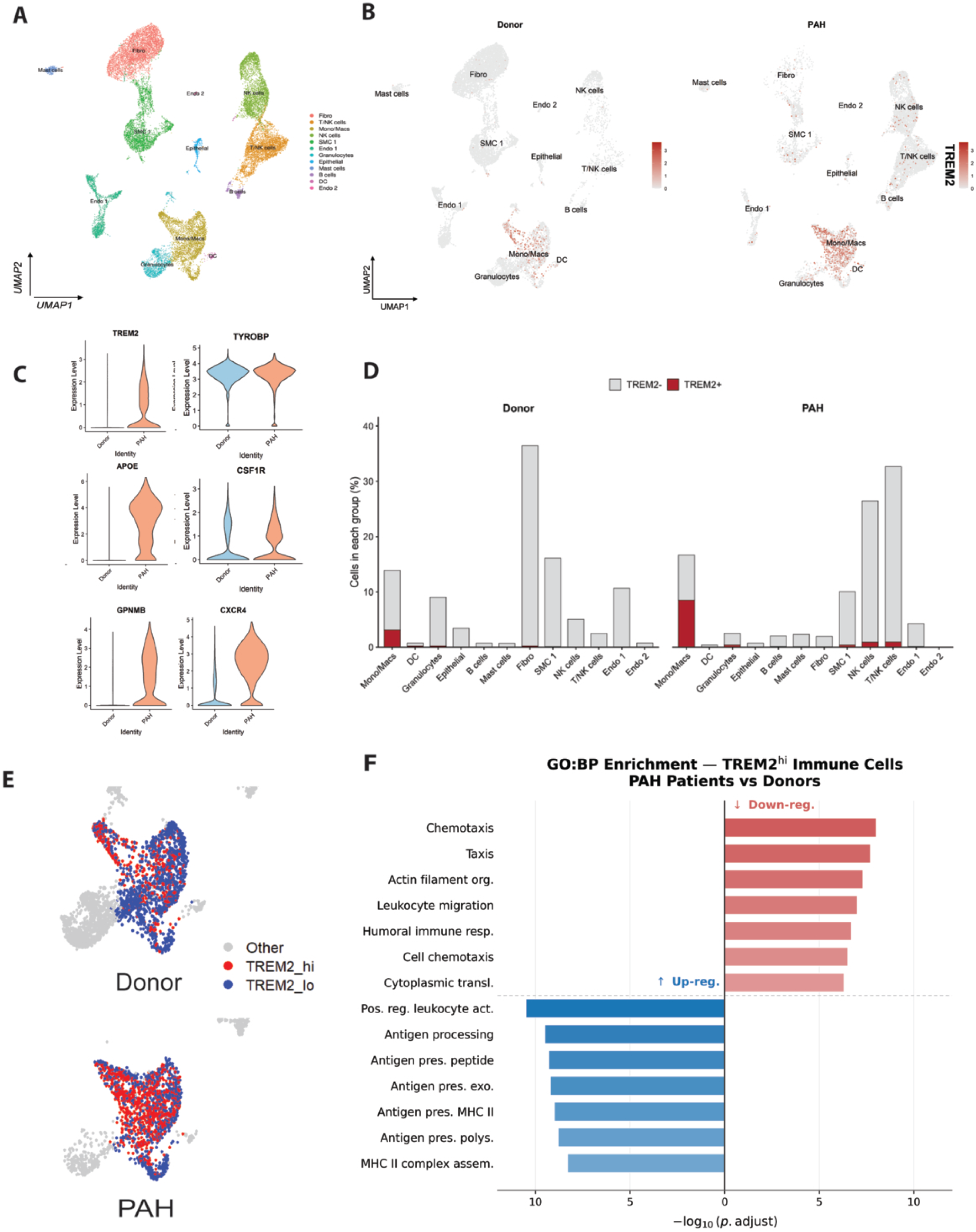
Patients with Pulmonary Arterial Hypertension (PAH) Exhibit High Levels of TREM2 Expression In Mono/macsPopulations in Pulmonary Arteries. A. UMAP visualization of publicly available scRNA-seq dataset GSE210248 derived from human pulmonary artery tissue (PAH patients, n=3; donors, n=3). All sequenced cells colored by annotated cell type. Twelve major populations were identified. B. UMAP feature plots overlaying TREM2 transcript expression onto all cells in donor (left) and PAH (right) tissue. Color scale indicates transcript expression level. C. UMAP plots of Mono/Macs cluster from donor (top) and PAH (bottom) tissue classified as TREM2-high (red), TREM2-low (blue), or other (grey). D. Bar plots showing relative contribution of each cell type to total TREM2 transcript expression in Donors versus PAH. E. Violin plots showing expression of TREM2, TYROBP, APOE, CSF1R, GPNMB, and CXCR4 in the Mono/Macs cluster in donors (left) and PAH patients (right). F. GO:BP enrichment analysis of differentially expressed genes in TREM2-high Mono/Macs from PAH patients versus donors. Red bars indicate significantly downregulated gene sets; blue bars indicate upregulated gene sets (−log₁₀ adjusted P value). DC indicates dendritic cells; GO:BP, Gene Ontology Biological Process; Mono/Macs, monocytes/macrophages; scRNA-seq, single-cell RNA sequencing; SMC, smooth muscle cells; TREM2, triggering receptor expressed on myeloid cells 2; TYROBP, TYRO protein tyrosine kinase-binding protein; UMAP, uniform manifold approximation and projection.

Visualization of canonical TREM2-associated and myeloid identity genes in the Mono/Macs cluster demonstrated supportive concomitant upregulation of APOE, GPNMB, and CSF1R in PAH compared to donors (Figure 1E). Additionally, CXCR4 expression was elevated in the PAH Mono/Macs cluster relative to donors (Figure 1C).

Mapping of TREM2 expression across multiple cell types suggested higher proportions of TREM2-positive cells between PAH and donor tissue, especially in mono/macs populations (Figure 1D). In healthy donor tissue, TREM2 expression was almost entirely restricted to the mono/macs population, with negligible contributions from other cell types, and the overall magnitude of TREM2 expression across the cell types was low. In PAH, the total magnitude of TREM2 expression across the tissue was markedly higher than in donors (Figure D, Figure S2), acknowledging that while mono/macs monocytes and macrophages remained the major contributor to total TREM2 expression, expression was also observed in other cell populations.

To further characterize TREM2 expression within the mono/macs compartment, cells classified as TREM2_hi (normalised expression ≥0.3) or TREM2_lo (normalised expression <0.3), following the threshold established by Cui et al.^23^ This analysis revealed a marked expansion of the TREM2_hi subpopulation in PAH relative to healthy donors (Figure 1E). Across the three samples in each group, 51% of mono/macs were classified as TREM2_hi in PAH, compared with 22.5% in donor controls (Figure S3). The proportion of TREM2-high monocytes/macrophages was higher in each PAH sample than in any donor sample, with mean sample-level proportions of 43.2% versus 23.7%. Furthermore, gene ontology analysis of differentially expressed genes that are up or downregulated in the TREM2^hi^ cluster identified two opposing transcriptional programs (Figure 1F). Gene sets for chemotaxis, actin filament organization, leukocyte migration, humoral immune response, cell chemotaxis, and cytoplasmic translation were significantly downregulated in PAH TREM2^hi^ cells. Conversely, positive regulation of leukocyte activation, antigen processing and presentation, peptide-MHC II complex-mediated antigen presentation, exogenous antigen presentation, polymorphic MHC II presentation, and MHC II complex assembly were significantly upregulated. This transcriptional profile is consistent with a metabolically active, tissue-resident macrophage phenotype with enhanced immunoregulatory and antigen-presenting functions. Together, these findings suggest that alterations in immune cell composition and responses in PAH may involve TREM2-associated myeloid activation, providing strong rationale for testing the role of TREM2 using *in vivo* models of disease.

### TREM2 deficiency protects against chronic hypoxia-induced pulmonary hypertension

To determine whether TREM2 contributes to the development of PH, wild-type (WT) and TREM2 knockout (TREM2 KO) mice, males and females, were exposed to normoxia (Nx, normobaric) or chronic hypoxia (Hx; FiO2 10%) for 28 days, and hemodynamic and structural indices of PH were assessed. Importantly, no significant difference was observed in hemodynamic outcomes between male and female mice (Figure S4).

TREM2 KO mice exposed to hypoxic conditions exhibited a significantly attenuated increase in RVSP compared to the control group. Related, TREM2 KO mice showed significantly lower maximum rate of right ventricular pressure rise (+dP/dt) compared to WT mice exposed to hypoxia, preventing the increase in contractile demand. Similarly, hypoxia altered the maximum rate of right ventricular pressure decline (−dP/dt) in WT mice, an effect that was abolished in TREM2-deficient mice (Figure 2A). Despite the reduction in pulmonary pressure, the Fulton index, measured by the RV/(LV+S) ratio, did not differ between hypoxic WT and TREM2 KO mice (Figure 2B).

**Figure 2.**
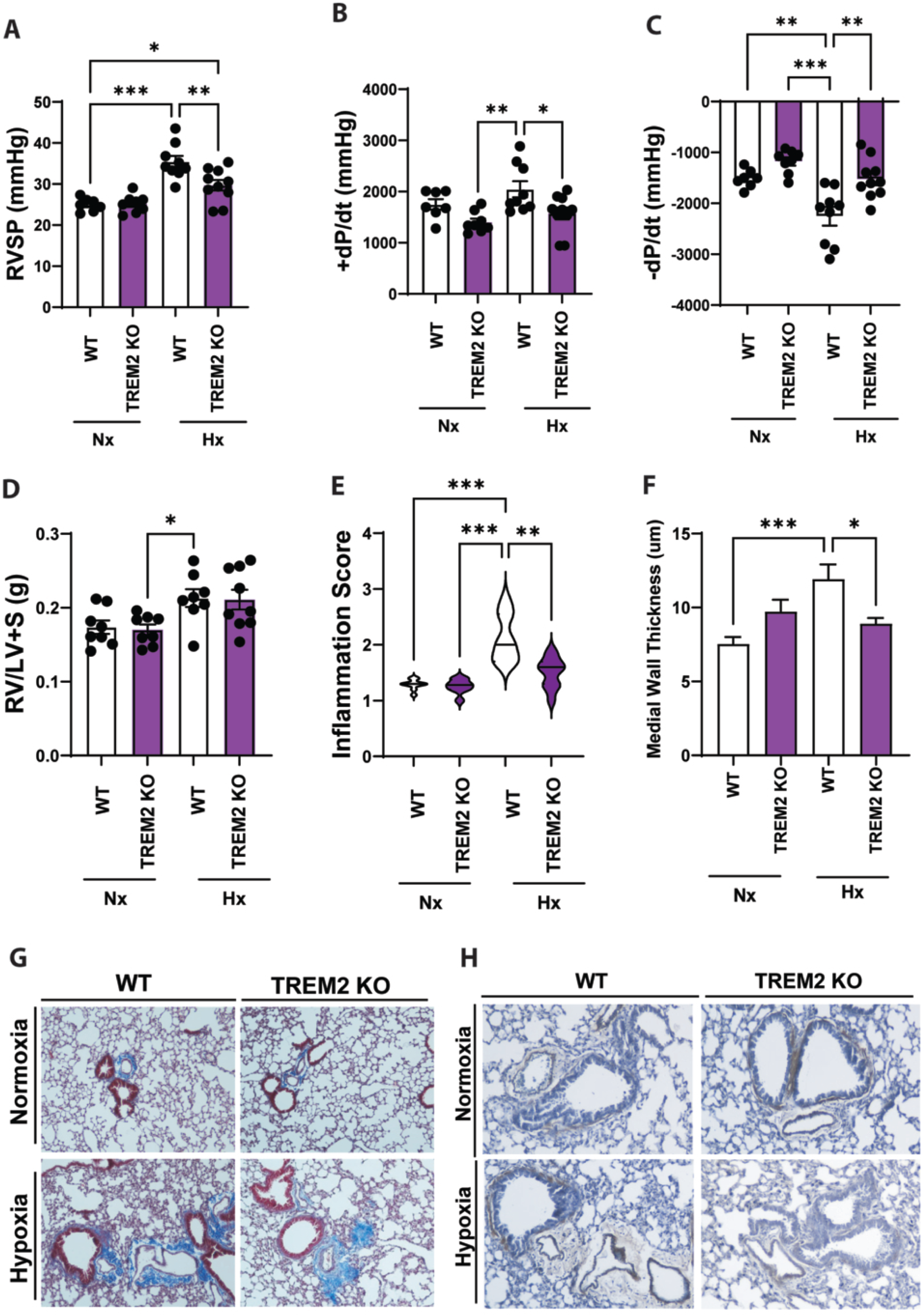
TREM2 Deficiency Leads to Significant Protection Against Hypoxia-Induced Pulmonary Hypertension. A. Hemodynamic and histological assessment of right ventricular and pulmonary changes in wild-type (WT) and TREM2 knockout (TREM2 KO) mice exposed to normoxia (nx) or hypoxia (hx) for 28 days; right ventricular systolic pressure (RVSP; left); B. Maximum rate of right ventricular pressure rise (+dp/dt; center). Maximum rate of right ventricular pressure decline (−dp/dt; right); B. Right ventricular hypertrophy index (RV/(LV+S)); C. Semi-quantitative pulmonary inflammation score on Masson trichrome–stained lung sections (scale 0–4; 10 fields per section, 10× magnification). C. Violin plots show median, interquartile range, and data distribution; D. Median pulmonary arterial wall thickness (μm) of small vessels; E. Representative Masson’s trichrome-stained lung sections showing vascular remodeling and collagen deposition (blue) in each group; F. Representative images of α-smooth muscle actin–stained pulmonary vessels illustrating muscularization under each condition. Statistical significance was determined by 2-way ANOVA with Tukey post-hoc correction (A, B and D) and Welch unpaired 2-tailed t test (C); P<0.05; **P<0.01; ***P<0.001. Hx indicates chronic hypoxia; KO, knockout; LV+S, left ventricle plus septum; Nx, normoxia; RV, right ventricle; RVSP, right ventricular systolic pressure; TREM2, triggering receptor expressed on myeloid cells 2; WT, wild-type; α-SMA, α-smooth muscle actin.

### TREM2 deletion reduces pulmonary inflammation and vascular remodeling

Given the human data demonstrating derangement in the inflammatory milieu associated with PAH, we next examined the preclinical model for evidence of alteration in inflammatory response due to TREM2 expression. Histological analysis demonstrated that chronic hypoxia significantly increased pulmonary inflammation in WT animals (p = 0.0021; Figure 2C) while TREM2 deficiency significantly attenuates this response (TREM2 KO-Hx vs WT-Hx, p = 0.0206). Representative images of Masson’s trichrome staining illustrate these findings showing marked perivascular fibrosis and structural remodeling in WT mice exposed to hypoxia (Figure 2E). In contrast, TREM2 KO mice exhibited visibly reduced inflammation and preservation of vascular architecture, consistent with attenuated fibrotic remodeling and the lower inflammation score (Figure 2E).

Consistent with these findings, α-Smooth muscle actin (α-SMA) staining demonstrated decreased pulmonary-vessel muscularization in hypoxic TREM2-deficient mice compared with hypoxic WT mice (Figure 2F). Quantitative assessment of vascular remodeling demonstrated that TREM2 deficiency prevented the increase in medial wall thickening in small pulmonary vessels induced by hypoxia, indicating protection from hypoxia-induced vascular hypertrophy (Figure 2D). Importantly, no significant hemodynamic or histological differences were observed between WT and TREM2 KO mice under normoxia. Together, these data indicate that TREM2 expression contributes to the development of chronic hypoxia-induced PH.

### TREM2 regulates mTOR activation, CCR2-mediated migratory priming, and T cell suppressive capacity of bone marrow MDSCs

Based on prior evidence that PH is associated with bone marrow myeloid dysregulation and that bone marrow-derived MDSCs contribute to pulmonary vascular remodeling,^26–28^ we examined whether TREM2 deficiency alters hypoxia-associated myeloid signaling within the bone marrow. Bone marrow cells from WT and TREM2 KO mice exposed to normoxia or hypoxia were analyzed by flow cytometry. Given that TREM2 signals through the PI3K/AKT/mTOR axis^23^ and that CCR2-dependent monocyte recruitment is a recognized feature of PAH progression^27,28^, we next examined mTOR, p-S6, and CCR2 expression in CD11b⁺ in the two major MDSC subsets: monocytic MDSCs (Mo-MDSCs) and polymorphonuclear MDSCs (PMN-MDSCs). In both subsets, hypoxia-induced phosphorylation of S6 (p-S6) was reduced in TREM2-deficient mice compared with hypoxic WT mice, indicating attenuated downstream mTOR signaling in the absence of TREM2 (Figure 3A). In Mo-MDSCs, hypoxia increased p-S6 expression in WT mice (p < 0.0001 vs. WT normoxia), whereas this response was significantly reduced in hypoxic TREM2 KO mice (p < 0.001 vs. hypoxic WT). Similarly, p-S6 expression was elevated in PMN-MDSCs from hypoxic WT mice (p < 0.0001 vs. WT normoxia) and attenuated in hypoxic TREM2 KO mice (p < 0.0001 vs. hypoxic WT). Total mTOR expression increased with hypoxia in Mo-MDSCs in both genotypes but was not significantly altered in PMN-MDSCs.

**Figure 3.**
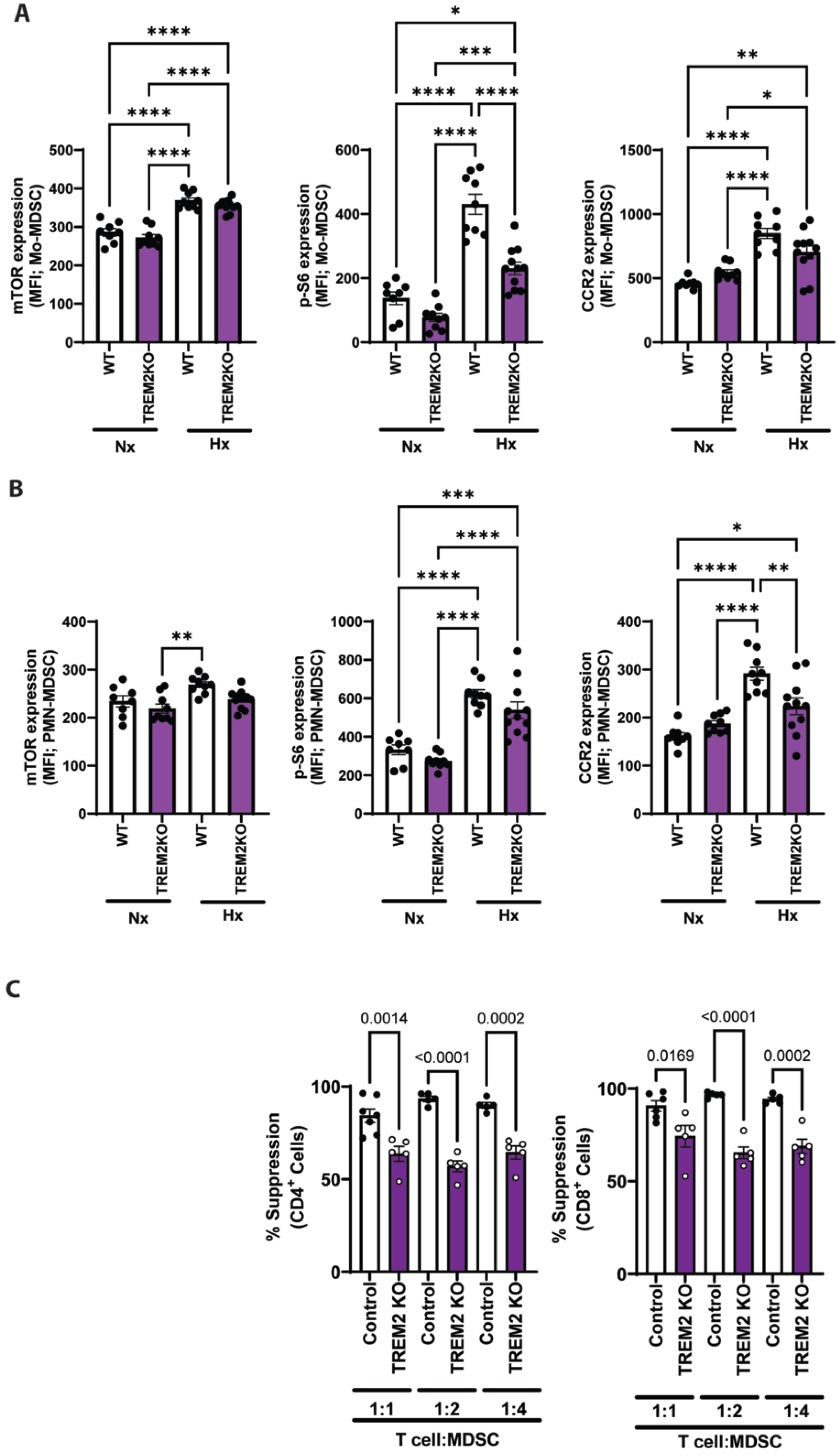
TREM2 Drives mTOR/p-S6 Activation and CCR2 Upregulation in Bone Marrow MDSCs and is Required for Full MDSC-Mediated T-Cell Suppression. WT and TREM2 KO mice were exposed to Nx or Hx for 28 days (n=6–10/group). A. Flow cytometric quantification of mTOR (left), p-S6 (center), and CCR2 (right) expression (MFI) in bone marrow Mo-MDSCs (CD11b⁺Ly6C^hi^). B. Flow cytometric quantification of mTOR (left), p-S6 (center), and CCR2 (right) expression (MFI) in bone marrow PMN-MDSCs (CD11b⁺Ly6G⁺Ly6C^lo^). C. Ex vivo T-cell suppression assay. CellTrace Violet–labeled splenic T cells from naïve WT mice were co-cultured with splenic MDSCs from control or TREM2 KO mice at T cell:MDSC ratios of 1:1, 1:2, and 1:4 for 5 days with anti-CD3/CD28 stimulation. Percent suppression of CD4⁺ (left) and CD8⁺ (right) T-cell proliferation quantified by CellTrace Violet dilution (n=5–6/group). Data are mean±SEM. Each dot represents an individual mouse. Statistical significance was determined by 2-way ANOVA with Tukey post-hoc correction. *P<0.05; **P<0.01; ***P<0.001; ****P<0.0001. CCR2 indicates C-C chemokine receptor type 2; Hx, chronic hypoxia; MDSC, myeloid-derived suppressor cell; MFI, mean fluorescence intensity; Mo-MDSC, monocytic MDSC; mTOR, mechanistic target of rapamycin; Nx, normoxia; PMN-MDSC, polymorphonuclear MDSC; p-S6, phosphorylated ribosomal protein S6; TREM2, triggering receptor expressed on myeloid cells 2; WT, wild-type.

Interestingly, hypoxia significantly increased cell-surface CCR2 expression in bone marrow Mo-MDSCs and PMN-MDSCs from WT mice. In PMN-MDSCs, this hypoxia-induced increase was attenuated in TREM2 KO mice compared to hypoxia WT (p < 0.01 vs. hypoxic WT; Figure 3), whereas CCR2 expression did not differ between hypoxic WT and TREM2 KO Mo-MDSCs. These data are consistent with evidence that CCL2-CCR2 signaling initiates downstream activation of PI3K/AKT/mTOR axis in myeloid cells ^29,30^ and together with our current findings with mTOR/p-S6, suggest that TREM2 may promote a hypoxia-responsive, CCR2-associated migratory phenotype specifically in PMN-MDSCs.

The suppressive capability of MDSCs is known to influence development of PH. We therefore hypothesized that TREM2 plays a role in the myelosuppressive capacity of these cells. To test this, we evaluated the capacity to suppress T cell proliferation, using a validated suppression assay, in which MDSCs isolated from control and TREM2 KO mice were co-cultured with activated T cells isolated from naïve WT control mice (Figure 3C). Consistent with the flow cytometric data, MDSCs from TREM2 KO mice demonstrated significantly reduced suppression of both CD4+ and CD8+ T cell proliferation compared to control MDSCs across all ratios tested (Figure 3C). Interestingly, in contrast to bone marrow-derived data, no significant differences in mTOR-signaling were observed between WT and TREM2-deficient mice myeloid cells isolated from the lung under hypoxic conditions (Figure 4). Moreover, CCR2 expression was increased in CD11b⁺ and PMN-MDSCs isolated from the lungs of TREM2-deficient hypoxic mice, highlighting this compartment-specific role of TREM2 in orchestrating myeloid cell mobilization in PH. In aggregate, however, these findings demonstrate that TREM2 expression is required for MDSC immunosuppressive function, a plausible contributing mechanism to PH.

**Figure 4.**
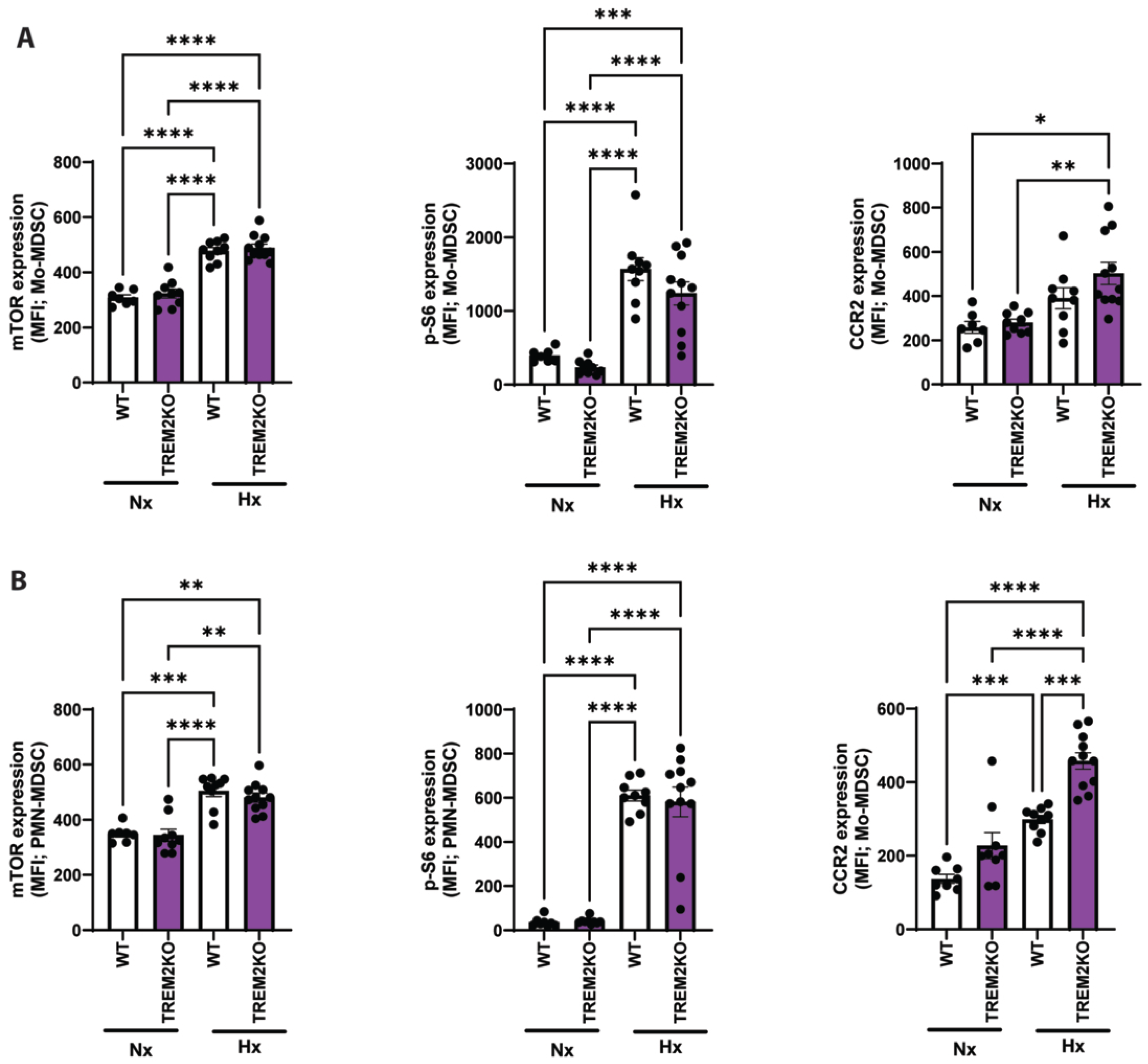
TREM2 Deficiency is Associated with Increased mTOR/p-S6 Activation and Elevated CCR2 Expression in Pulmonary Myeloid Populations Under Hypoxia. WT and TREM2 KO mice were exposed to Nx or Hx for 28 days (n=6–10/group). Top row: flow cytometric quantification of mTOR (left), p-S6 (center), and CCR2 (right) expression (MFI) in lung Mo-MDSCs (CD11b⁺Ly6C^hi^). Bottom row: flow cytometric quantification of mTOR (left), p-S6 (center), and CCR2 (right) expression (MFI) in lung PMN-MDSCs (CD11b⁺Ly6G⁺Ly6C^lo^). Data are mean±SEM. Each dot represents an individual mouse. Statistical significance was determined by 2-way ANOVA with Tukey post-hoc correction. *P<0.05; **P<0.01; ***P<0.001; ****P<0.0001. CCR2 indicates C-C chemokine receptor type 2; Hx, chronic hypoxia; MDSC, myeloid-derived suppressor cell; MFI, mean fluorescence intensity; Mo-MDSC, monocytic MDSC; mTOR, mechanistic target of rapamycin; Nx, normoxia; PMN-MDSC, polymorphonuclear MDSC; p-S6, phosphorylated ribosomal protein S6; TREM2, triggering receptor expressed on myeloid cells 2; WT, wild-type.

### TREM2 expression is increased in circulating myeloid-derived suppressor cells and correlates with disease severity in pulmonary hypertension

To validate the clinical relevance of our findings in patients with PH, we assessed TREM2 protein levels in circulating myeloid populations and whether they associate with indices of disease severity. For that purpose, we performed flow cytometric analysis of peripheral blood mononuclear cell from PAH patients and control donors (Figure 5A, Table 1). Consistent with our preclinical findings quantification of TREM2 expression within circulating MDSCs revealed significantly higher levels in PAH subjects compared to controls (Figure 5B; P < 0.01).

**Figure 5.**
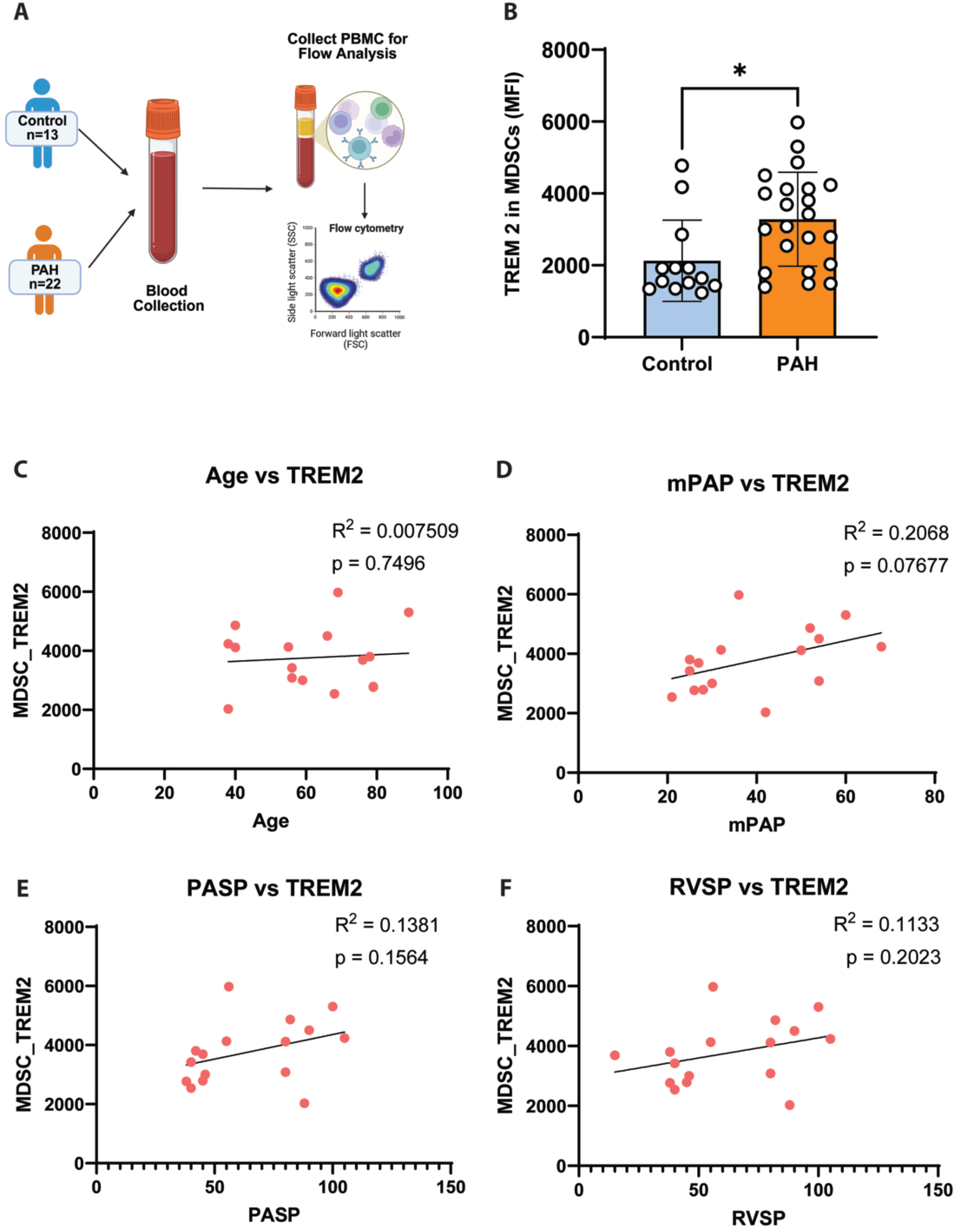
TREM2 Expression is Elevated in Circulating MDSCs from PAH Patients and Shows a Directional Association With Hemodynamic Severity. Blood samples were collected from healthy controls (n=13) and PAH patients (n=22), and PBMCs were isolated and differentiated with GM-CSF for 7 days before flow cytometric analysis. A. Schematic representation of the study design. Peripheral blood mononuclear cells (PBMCs) were isolated and analyzed by flow cytometry to assess TREM2 expression in myeloid-derived suppressor cells (MDSCs). B. Quantification of TREM2 expression (mean fluorescence intensity, MFI) in circulating MDSCs from healthy control and PAH patients. Each dot represents an individual subject. Statistical significance was determined by unpaired Welch t test. *P<0.05. C–F. Correlation analyses between TREM2 MFI expression in MDSCs and clinical parameters, including age (C); mean pulmonary arterial pressure (mPAP) (D), pulmonary artery systolic pressure (PASP) (E), and right ventricular systolic pressure (RVSP) (F). GM-CSF, granulocyte-macrophage colony-stimulating factor; MDSC, myeloid-derived suppressor cell; MFI, mean fluorescence intensity; mPAP, mean pulmonary arterial pressure; PAH, pulmonary arterial hypertension; PASP, pulmonary arterial systolic pressure; PBMC, peripheral blood mononuclear cell; R², coefficient of determination; RVSP, right ventricular systolic pressure; SSC, side light scatter; TREM2, triggering receptor expressed on myeloid cells 2.

**Table 1:**
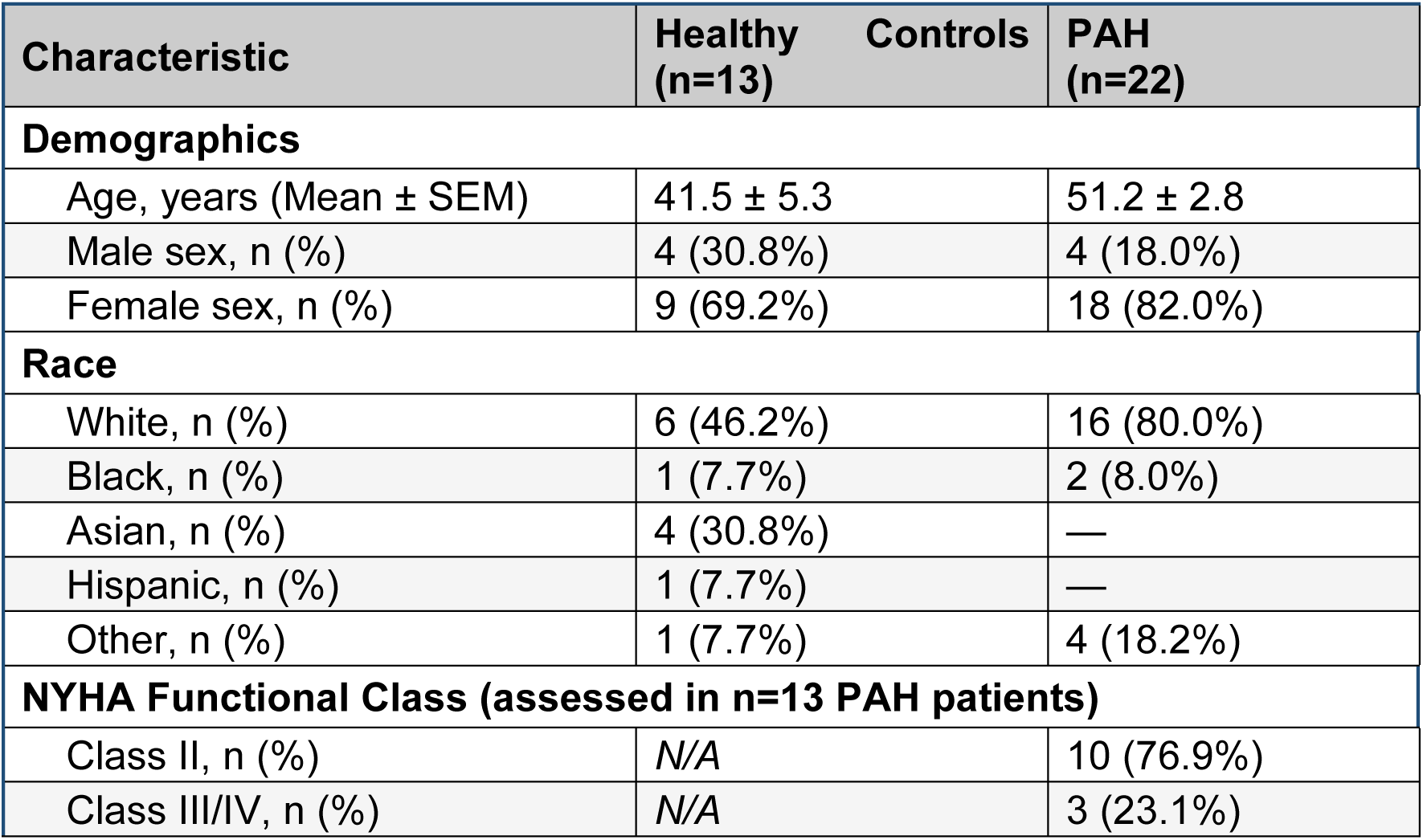
Baseline Demographic and Clinical Characteristics. PAH, pulmonary arterial hypertension; NYHA, New York Heart Association; SEM, standard error of the mean; N/A, not applicable; —, not available. Data presented as mean ± SEM or n (%).

We next examined for correlation between TREM2 expression and patient outcomes and demographic features. First, we confirmed that in our cohort of patients TREM2 levels were not simply altered due to aging (Figure 5C). Interestingly, TREM2 expression levels in circulating myeloid-derived suppressor cells showed a positive correlation with mean pulmonary arterial pressure (mPAP; R² = 0.20, p = 0.07), confirming an association between TREM2 upregulation and hemodynamic findings at presentation. Taken together, these findings suggest a modest but consistent directional association between TREM2 expression in circulating MDSCs and hemodynamic indices of disease severity in PH (Figure 5D–E). While hypothesis-generating given patient numbers, the consistency and directionality of the associations across hemodynamic parameters strengthen the potential role of TREM2 as a therapeutic target and biomarker in PH.

## Discussion

MDSCs have been increasingly implicated in PH-associated vascular remodeling. These cells expand in the bone marrow and can accumulate in peripheral tissues and regulate inflammatory and adaptive immune responses.^29^ Prior work from our group and others has identified increased MDSC accumulation in experimental PH and in patients with PH.^25,31,32^ Here, we identify TREM2 as a potential regulator of the pathogenic and immunosuppressive functions of MDSCs in PH. Using complementary approaches spanning human transcriptomic analysis, genetic loss-of-function experiments, and clinical correlative data, we demonstrate that: (1) TREM2 is selectively upregulated in mono/macs populations in PAH pulmonary arteries, accompanied by a disease-associated myeloid transcriptional program; (2) global TREM2 deficiency attenuates CH-PH, right ventricular dysfunction, pulmonary inflammation, and vascular remodeling in mice; (3) TREM2 deficiency was associated with reduced mTOR/p-S6 activation and CCR2 abundance in bone marrow MDSCs which is required for their T cell suppressive function; and (4) TREM2 is elevated in circulating MDSCs from PAH patients and shows a directional association with hemodynamic severity. Together, these findings support TREM2 as a central component of the immunosuppressive niche that drives PAH progression.

The scRNA-seq analysis reveals that TREM2 upregulation in the PAH pulmonary vasculature is not a generalized inflammatory response but a program centered on the Mono/Macs compartment, with coordinate induction of DAP12/TYROBP, APOE, CSF1R, and GPNMB, the canonical signature of disease-associated macrophages (DAMs) described in neurodegeneration, non-alcoholic steatohepatitis, and, most recently, pulmonary fibrosis.^23,24,33–35^ The restriction of TREM2 expression to Mono/Macs in donor tissue, contrasted with its broader distribution across multiple immune and structural cell populations in PAH, suggests that the hypoxic, lipid-enriched, and DAMP-rich PAH vascular microenvironment may induce TREM2 expression in cell types that do not constitutively express it. This is consistent with evidence that TREM2 is activated by oxidized phospholipids, ApoE, and DAMPs that accumulate in chronically injured tissues. ^19,33,36–38^ For example, in a COVID-19 cohort, TREM2 surface expression was “hardly detected” on CD4+ or CD8+ T cells from healthy donors, but was strongly induced in circulating and lung-infiltrating T cells from patients^39^. While a well-recognized limitation in scRNA-seq is that unsupervised clustering does not yield perfectly pure populations, these data nonetheless provide the first evidence that TREM2 contributes to PH, and future studies should consider the potential contribution of TREM2 across other cell types.

A key observation in this study is that the hemodynamic protection observed in TREM2 KO mice, with reduced RVSP and attenuated right ventricular contractility, changes alongside reduced pulmonary inflammation and vascular remodeling, supporting the concept that TREM2 acts as a functional inflammatory “rheostat” in PH pathogenesis. The finding that TREM2 KO mice do not exhibit reduced right ventricular hypertrophy despite attenuated pressure elevation is notable and may reflect early structural right ventricular remodeling driven by mechanisms independent of TREM2, or a threshold effect in which 28 days of even modestly elevated afterload may initiate hypertrophic remodeling regardless of genotype. This dissociation between pressure and hypertrophy has been described in other protective models of hypoxic PH and warrants further investigation of factors beyond the magnitude of pressure elevation, including direct effects of hypoxia on the myocardium.^40,41^ These direct myocardial effects will need to be elucidated in the future.

TREM2 has been studied most extensively in neurodegenerative disease, where it has been linked to APOE. APOE serves both as a ligand for TREM2 and as a key transcriptional target induced by TREM2 signaling, a program that regulates lipid handling, cellular survival, and disease-associated myeloid-state transitions.^21,42,43^ In these contexts, TREM2-associated signaling has also been linked to downstream metabolic pathways, including PI3K/AKT/mTOR signaling, which may support myeloid-cell adaptation to chronically injured tissue microenvironments. In our scRNA-seq analysis, APOE was increased within the PAH Mono/Mac compartment. This finding contrasts with earlier reports of reduced APOE expression in whole lung tissue from patients with PAH and with studies identifying APOE as an antiproliferative mediator in pulmonary artery smooth muscle cells downstream of BMP-2/PPARγ signaling.^44^ The apparent discrepancy may reflect cell type-specific functions of APOE in pulmonary vascular disease: whereas reduced APOE in structural vascular cells may favor proliferative remodeling, increased APOE within myeloid cells may identify a distinct TREM2-associated immunometabolic state. This mirrors a compartment-dependent divergence we previously described for CXCR2, which is causative for PH when expressed by myeloid cells but protective when expressed by endothelium. ^45^ Therefore, although future studies are still required to fully understand the APOE contributions to PH, Our data lay the foundational framework for connecting known changes in APOE in PH, innate immune activity, and metabolic signaling.

The transcriptional profile of TREM2^hi^ PAH macrophages in our analysis demonstrated reduced expression of gene programs related to chemotaxis, actin dynamics, and leukocyte migration alongside upregulation of antigen processing and MHC II presentation. This transcriptional profile is consistent with a post-migratory, tissue-committed macrophage identity recently described in pulmonary fibrosis, where TREM2 upregulation silence CCR2 upon tissue entry and adopt a sessile, immunosuppressive phenotype sustained by sphingolipid ligand sensing and CCL2-dependent myeloid recruitment.^24^ The competition rather than absence of this recruitment process is directly supported by our preclinical data, in which TREM2 KO lung myeloid cells retain elevated CCR2 expression under hypoxia, consistent with failure to complete TREM2-dependent post-migratory differentiation. The convergence of lipid ligand sensing, CCR2-dependent recruitment, and immunosuppressive tissue macrophage identity across both fibrotic and hypertensive pulmonary disease positions TREM2 as a conserved pathological macrophage state that operates across chronic lung disease contexts, shaped by the specific microenvironmental cues of each disease niche.

Notably, TREM2 deficiency was associated with distinct effects on CCR2 expression in bone marrow and lung PMN-MDSCs. In the bone marrow, TREM2 attenuates the hypoxic upregulation of CCR2, whereas in the lung it potentiates that same induction. This indicates that TREM2 signaling supports induction of the CCR2⁺ migratory program during myeloid development in the marrow. However, in the lungs, cells from TREM2 KO mice fail to complete the maturation program that downregulate CCR2.

The opposing effects of TREM2 we observe across compartments are consistent with its broader biology, in which TREM2 plays context-dependent and sometimes opposing roles depending on tissue and disease development^35,36,46^. While in some conditions such as neurodegenerative disease TREM2 could be protective, in cancer, TREM2 marks immunosuppressive tumor-associated myeloid cells, and its deletion or antibody blockade relieves this suppression and enhances anti-tumor T-cell responses^35^.

Therefore, in our next step, the *ex vivo* T cell suppression assay provided direct functional evidence that TREM2 is required for the full immunosuppressive capacity of MDSCs. TREM2 KO MDSCs suppressed both CD4⁺ and CD8⁺ T cell proliferation significantly less effectively than WT MDSCs across the tested MDSC: T-cell ratios, in a dose-dependent manner. This finding has been implicated in pulmonary vascular inflammation and remodeling in PAH^9^ and provide a potential mechanism whereby TREM2 KO mice are protected against CH-PH.

It is important to highlight that the direction and magnitude of TREM2’s effect are known to depend on disease stage and duration. In Alzheimer’s disease, TREM2 is protective during the early amyloid-driven phase but can become detrimental in later tau-driven neurodegeneration, and chronic TREM2 activation has been shown to exacerbate pathology.^22,47^ A recurring interpretation is that transient TREM2-driven responses support clearance and repair, whereas sustained activation may become maladaptive. Our findings are consistent with a severity-dependent scaling of this program. The chronic hypoxia model used here produces only mild PH and a correspondingly modest TREM2 signature, whereas our single-cell data from patients with severe pulmonary arterial hypertension (Figure 1) show greater TREM2 upregulation accompanied by more pronounced transcriptional changes in TREM2^hi^ immune cells, including downregulation of chemotaxis and migratory programs. Although cross-species and severity differences preclude a direct comparison, these observations are consistent with the possibility that the functional footprint of the TREM2 program intensifies with disease severity and may therefore be more pronounced in advanced human disease than in our mild preclinical model.

## CONCLUSION

The present study identifies TREM2 as a novel and functionally important regulator of myeloid-driven immunosuppression and vascular remodeling in PAH. TREM2 upregulation in PAH pulmonary artery macrophages defines a post-migratory, immunosuppressive myeloid identity that drives T cell suppression and vascular remodeling through mTOR-dependent metabolic activation and CCR2-regulated myeloid trafficking. TREM2 deficiency attenuates these effects, protecting against hypoxia-induced PH in mice while preserving T cell homeostasis. These findings establish TREM2 as a potential candidate for therapeutic targeting and biomarker development in PAH.

### Clinical/Pathophysiological Implications

Although in a modest cohort, our data provides the first translational evidence that the TREM2-associated myeloid program may contribute to PH. Correlative analysis with hemodynamic data is still preliminary, but sufficient to support future evaluation of TREM2 in larger prospective PH cohorts. From a therapeutic standpoint, anti-TREM2 strategies have recently demonstrated efficacy in preclinical pulmonary fibrosis models,^37,48,49^ and the present data provide a mechanistic basis for investigating similar approaches in PH.

### Limitations

The chronic hypoxia model produces relatively mild and reversible PH and does not fully reproduce the complex vascular lesions of human PAH. The human clinical cohort was modest in size (n=22 PAH, n=13 controls), and the association between circulating MDSC TREM2 expression and hemodynamic severity did not reach statistical significance. Future studies using conditional knockout models, larger clinical cohorts with longitudinal follow-up, and mechanistic dissection of TREM2 cell-specific contribution in the context of PAH will be important to fully define the therapeutic potential of TREM2 targeting in this disease.

### Novelty and Significance

- This study identifies TREM2 as a novel regulator of myeloid-driven immunosuppression and pulmonary vascular remodeling in PH;
- TREM2 acts upstream of multiple immunoregulatory effector mechanisms simultaneously: mTOR activation, CCR2-dependent trafficking, and T cell suppression, positioning TREM2 as a potential therapeutic target in PH;
- PH leads to significant upregulation of TREM2 expression in circulating myeloid cells, providing clinical evidence for TREM2 as a candidate for a non-invasive biomarker of myeloid disease activity, warranting prospective multicenter validation.

## Author contributions

A.C. Oliveira and A.J. Bryant designed the experiments, performed data analysis and interpreted results of experiments. A.C. Oliveira wrote the manuscript and prepared figures. F. Silva, generated the graphical abstract, edited and revised the manuscript. A.C. Oliveira, F. Silva, M.D. Alves, A.T. Pham, J. Alvarez-Castanon, S. Khanfar, R. Abdelnor, C. Phillips, M.T. Hall, Chunhua Fu, performed experiments. S.T. Vir and K.E. Ray, analyzed patient database with clinical and demographic data. S.T. Vir prepared the demographic table. Y. Zhang, C. Harris, and L. Chen analyzed scRNAseq data. M.D. Alves, performed histopathological analysis and scoring. A. de Kloet and E. Krause, A.J. Bryant edited and revised the manuscript. A.J. Bryant supervised the project. All authors approved the final version of the manuscript.

